# A highly conserved sRNA downregulates multiple genes, including a σ^54^ transcriptional activator, in the virulence mode of *Bordetella pertussis*

**DOI:** 10.1101/2024.11.19.624354

**Authors:** Minji Sim, Jeffers Nguyen, Karolína Škopová, Kyungyoon Yoo, Chin-Hsien Tai, Leslie Knipling, Qing Chen, David Kim, Summer Nolan, Rawan Elaksher, Nadim Majdalani, Hernan Lorenzi, Scott Stibitz, Kyung Moon, Deborah M. Hinton

## Abstract

Bacterial sRNAs together with the RNA chaperone Hfq post-transcriptionally regulate gene expression by affecting ribosome binding or mRNA stability. In the human pathogen *Bordetella pertussis*, the causative agent of whooping cough, hundreds of sRNAs have been identified, but their roles in *B. pertussis* biology are mostly unknown. Here we characterize a Hfq-dependent sRNA (S17), whose level is dramatically higher in the virulence (Bvg^+^) mode. We show that transcription from a σ^A^-dependent promoter yields a long form of 190 nucleotides (nts) that is processed by RNase E to generate a shorter, more stable form (S17S) of 67 nts. Using RNA-seq and RT-qPCR, we identify 92 genes whose expression significantly increases in the absence of S17. Of these genes, 70 contain sequences at/near their ribosome binding sites (RBSs) that are complementary to single-stranded (ss) regions (Sites 1 or 2) of S17S. The identified genes include those encoding multiple transporters and 3 transcriptional regulators. Using a *lacZ* translational reporter system, we demonstrate that S17S directly represses one of these genes, *BP2158*, a σ^54^- dependent transcriptional regulator, suggesting the repression of a σ^54^ regulon in the Bvg^+^ mode. We find that the S17S region containing Sites 1 and 2 is 100% conserved throughout various Betaproteobacteria species, and the S17S target sites are often conserved in the homologs of the *B. pertussis* target genes. We speculate that S17S regulation represents a highly conserved process that fine-tunes gene expression in the Bvg^+^ mode of *B. pertussis* and perhaps under other conditions in related bacteria.

**IMPORTANCE:** Regulation of gene expression involves controlling transcription, translation, and transcript degradation. sRNAs with short sequences complementary to an mRNA sequence are involved in post-transcriptional regulation by aiding or interfering with either ribosome binding or nuclease attack. In the human pathogen *Bordetella pertussis*, the causative agent of whooping cough, hundreds of sRNA have been identified, but their functions remain largely unknown. We have characterized a sRNA that is abundant in the virulence mode of *B. pertussis* and serves to down-regulate multiple genes including transcriptional regulators and various transporters. We demonstrate that this sRNA directly represses a transcriptional factor, suggesting that it influences the regulation of specific *B. pertussis* regulons. The high conservation of this sRNA and its targets within Betaproteobacteria suggests a conserved pathway for gene regulation.

## INTRODUCTION

In all organisms, the ability of small non-coding RNAs (sRNAs) to anneal to mRNAs allows them to function as post-transcriptional regulators, modulating translation and mRNA stability (1–3). In bacteria, this process, which is often mediated by the RNA chaperone Hfq, regulates multiple pathways, providing a mechanism for adaption to various environmental stressors, including oxidative stress, DNA damage, iron and nutrition starvation, and lower temperatures (2, 4–8). For pathogenic bacteria, sRNA/Hfq regulation has been shown to affect virulence, by modulating myriad virulence determinants, such as those involved in quorum sensing, Type III secretion systems (T3SS), iron transport, and/or biofilm formation (9–14). The importance of *hfq* for this regulation is demonstrated by the changes in pathogenesis observed with *Δhfq* strains [reviewed in (14, 15)].

The obligate human pathogen *Bordetella pertussis* (*B. pertussis*) is a Gram-negative bacterium that causes whooping cough (pertussis), a highly contagious, acute respiratory illness (16). Although vaccination coverage for *B. pertussis* has been high for decades (∼86% worldwide according to WHO), the relatively short duration of immunity provided by the acellular vaccine that is offered in many developed countries, including the US, is thought to be involved in regional outbreaks and infant mortality (17).

Regulation of gene expression in *B. pertussis* arises in large part through the action of two-component systems that respond to changing conditions as the bacterium is expelled from one human respiratory tract and then travels into and through another. Virulence in *B. pertussis* is primarily controlled by a two-component system: BvgS, the histidine sensor kinase, and BvgA, the response regulator (18). Under standard laboratory growth at 37° C, phosphorylated BvgA (BvgA∼P), which is phosphorylated via a BvgS-mediated phosphorelay, predominates, resulting in the Bvg^+^ mode. In this mode, BvgA∼P dimers activate transcription of virulence-activated genes (*vags*), such as pertussis toxin (*ptx*), filamentous hemagglutinin/adhesion (*fha*), fimbriae (*fim3, fim2*), and *bvgR* (19–22). The *bvgR* gene encodes an EAL domain, typical of cyclic di-GMP (c-di-GMP) phosphodiesterases, which reduces the levels of c-di-GMP. This in turn negatively affects the activity of another RR, RisA, which requires c-di-GMP for the activation of a set of virulence-repressed genes (*vrgs*) (23). Consequently, in the Bvg^+^ mode, the activation of *bvgR* decreases the concentration of c-di-GMP, leading to *vrg* repression.

Although standard laboratory growth conditions result in the Bvg^+^ mode, lower temperature (25° C) or addition of sufficient concentrations of compounds, such as nicotinic acid or magnesium sulfate (MgSO_4_), modulates BvgS activity. Under these conditions, BvgA is not phosphorylated, *vags* and *bvgR* are not expressed, the concentration of c-di-GMP is high, RisA is active, and *vrgs* are expressed (24). This is the Bvg^-^ mode, which is thought to be important for transmission/persistence (25–27) and may also play a role in the macrophage environment (28). A third Bvg mode, Bvg^i^, represents a state with an intermediate level of BvgA∼P. Only one gene, *bipA*, is classified as Bvg^i^ since it is activated by levels of BvgA∼P that activate early *vags*, but it is repressed by the higher levels required for late gene activation (29). This state may be relevant during infection (30, 31).

Despite the global transcriptional regulation imposed by the response regulators BvgA in the Bvg^+^ mode and for RisA in the Bvg^-^ mode, recent evidence has indicated that *B. pertussis*, like many other bacteria, also relies on sRNAs to fine-tune gene expression through post-transcriptional regulation. Previous work has shown that a deletion of *hfq* in the clinical strain *B. pertussis* 18323 attenuates the expression of various virulence genes, including those within the T3SS (32). In *B. pertussis* Tohama I, the sRNA RgtA (33), whose expression is enhanced in the Bvg^-^ mode and which interacts directly with Hfq (34), targets genes involved in glutamate metabolism (35). Other work has identified several sRNAs present during growth or colonization of mouse trachea by *B. pertussis* 18323 (36). One of these, Bpr4, functions in the Bvg^+^ mode, up-regulating *fha* (adhesin filamentous hemagglutinin) during tracheal colonization (37).

Using the prokaryotic sRNA search program, ANNOgesic, we previously conducted a genome-wide transcriptomic search for Bvg^+^, Bvg^-^, and Bvg-independent sRNAs present in *B. pertussis* Tohama I (34). We predicted 143 possible candidates, of which 25 were Bvg^+^-specific and 53 were Bvg^-^-specific, and we confirmed 10 of these by Northern Blot analyses. One of these, designated S17, interacts directly with Hfq and is highly enriched in the Bvg^+^ mode.

Here we show that S17 exists in 2 forms: a longer 190 nucleotide (nt) product (S17L), whose transcription starts from a σ^A^-RNA polymerase (σ^A^-RNAP)-dependent promoter (P_S17_) and terminates with a factor-independent stem-loop terminator, and a predominant, shorter ∼70 nt RNA (S17S) that arises from processing of S17L by RNase E. To understand the role of S17, we performed RNA-seq and RT-qPCR to compare *B. pertussis* gene expression in a ΔS17 strain relative to wild type (WT). We identify 91 genes that are significantly upregulated in the ΔS17 strain. Of these, 70 genes have sequences within the 5’ untranslated regions (UTRs) near or overlapping their ribosome binding sites (RBS) that are complementary to predicted single-stranded segments of S17S. Using a reporter system, we show that overexpression of S17S down-regulates translation of the predicted σ^54^-transcriptional regulator encoded by *BP2158*, a dehydrogenase (encoded by *mmsB*), and an operon encoding carbohydrate-binding ABC transporter genes (*BP2642A, BP2642, BP2641, BP2640, BP3638*). Furthermore, for the transcriptional regulator, we show that mutations affecting the putative annealing site disrupts regulation while complementary compensatory mutations restore regulation. Our results suggest that S17 together with Hfq serves to post-transcriptionally down-regulate sets of genes in the Bvg^+^ mode of *B. pertussis*.

## RESULTS

### Characterization of the *B. pertussis* sRNA S17

Our previous work searching for *B. pertussis* sRNAs (34) identified the intergenic sRNA S17, located between *BP3151*, encoding a glycosyltransferase, and *BP3152* (*coq7*), encoding 2- polyprenyl-3-methyl-6-methoxy-1,4-benzoquinone monooxygenase (Fig. 1A). The entire sequence and the genomic position of S17 is highly conserved among *B. pertussis*, *B. parapertussis,* and *B. bronchiseptica* species (34). Northern analyses indicated that S17 consists of 2 species: a low abundance, longer form S17L of ∼ 190 nt and a much more abundant, shorter form S17S of ∼70 nts [Fig. 1B; (34)].

**FIG 1.**
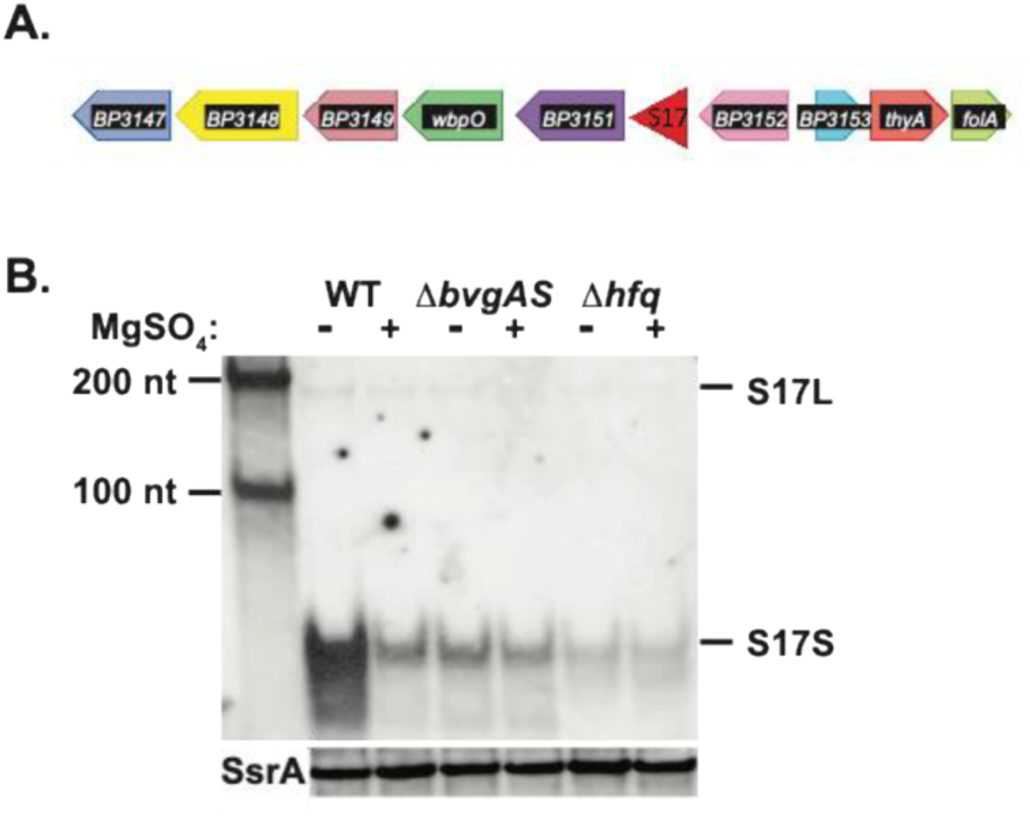
Genomic location and Bvg^+^-mode and Hfq enhancement of S17. (A) The position of S17 within the *B. pertussis* Tohama I genome is indicated. (B) Representative Northern blot (one of 2 biological replicates) shows the presence of S17 isolated from the indicated *B. pertussis* strains (WT, Δ*bvgAS*, or Δ*hfq*) grown in the absence or presence of MgSO_4_. First lane contains RNA size markers of the indicated lengths. S17L and S17S are marked; a rehybridization of the gel using a probe for the control sRNA SsrA is shown underneath.

To identify the 5’-end of S17, we used 5’-RACE analyses (5’-rapid amplification of cDNA ends), which revealed an ‘A’ 5’-end for S17L, located 7 bp downstream of a potential σ^A^- dependent promoter (Fig. 2A). The sequence of the putative promoter (P_S17_) contains good matches to the ideal −10 and −35 elements (5’TACTAT3’ and 5’TTGCAT3’ *vs.* consensus sequences of 5’TAtaaT3’ and 5’TTGaca3’, respectively) for σ^A^-RNAP, separated by the ideal spacer length of 17 bp. Transcription initiating here and ending at a predicted factor-independent terminator, a strong stem-loop followed by a run of 9 T’s, would generate an RNA of 190 nt to the end of the T-run, consistent with the size of S17L observed by the Northern analysis.

**FIG 2.**
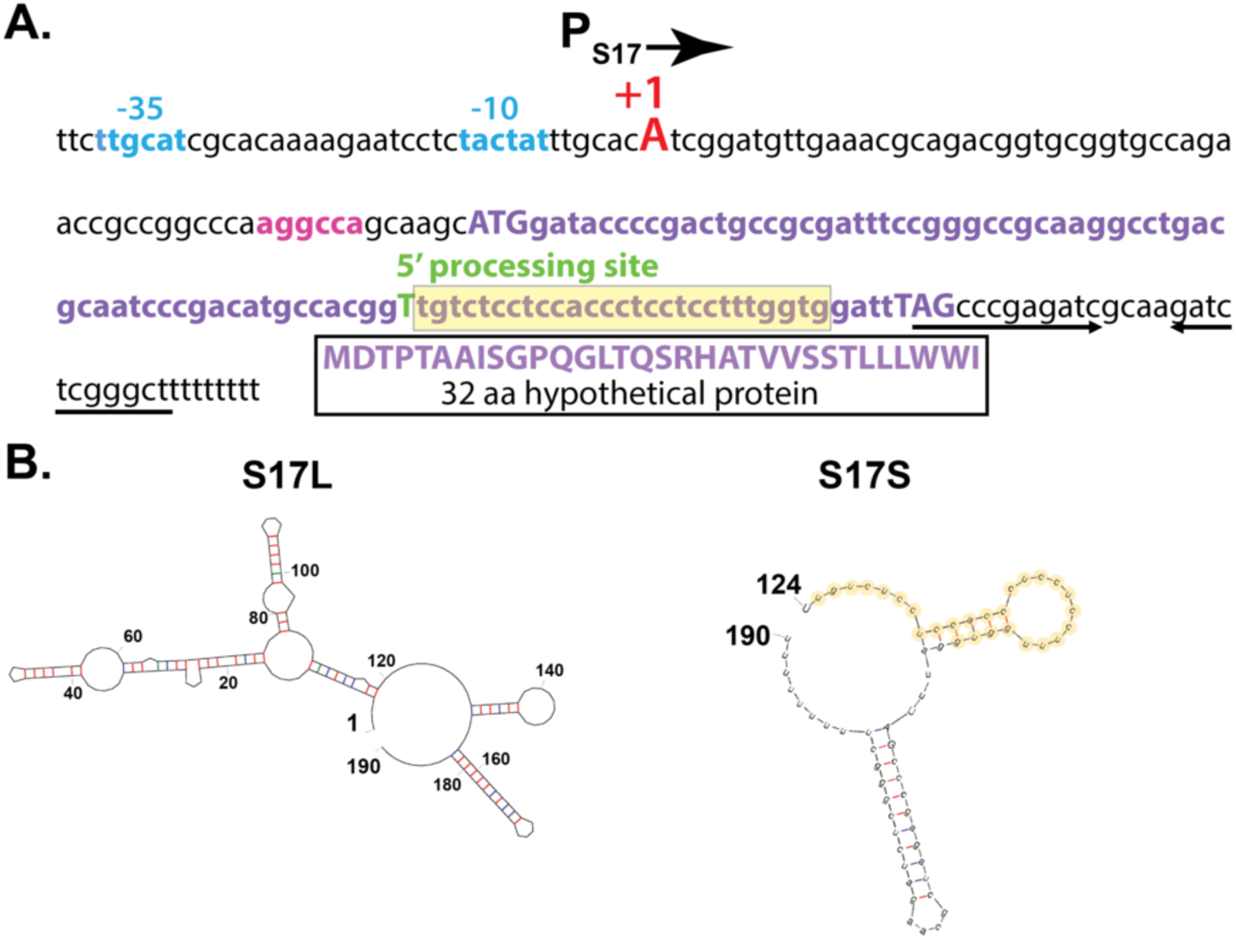
DNA sequence surrounding S17 and the predicted secondary structures of S17L and S17S RNAs. (A) S17 DNA sequence is shown as follows: promoter P_S17_ (assigned −35 and −10 elements in Carolina blue, +1 TSS for S17L capitalized in red); 5’UTR upstream of the small ORF with the position of the predicted RBS highlighted in magenta; the small ORF in purple (with the ATG start codon and the TAG stop codon capitalized) except for the 5’-processed end of S17S which is capitalized and shown in green; and the 3’UTR region with the factor independent stem-loop terminator shown designated by the arrows. The translated sequence of the 32 aa predicted protein is indicated. (B) Predicted mfold (74) secondary structures for S17L (left) and the processed S17S (right) are shown. Nucleotide positions relative to the +1 TSS are indicated. G:C, A:U, and G:U bps are shown by the red, blue, and green lines, respectively. The nucleotides highlighted in yellow in S17S in (A) and (B) represent the region that is 100% conserved in various Betaproteobacteria species (see Fig. 3).

5’-RACE analysis identified the 5’-end of S17S as a downstream T (Fig. 2A), which would generate a 67 nt RNA, consistent with the estimated size of S17S from the Northern blot. No identifiable promoter sequence was positioned upstream, suggesting that S17S might arise from processing. The 5’-end of S17S lies near the start of a C/U-rich loop that is located between putative stem loops. Our previous work indicated that the entire sequence of S17 is highly conserved in *B. pertussis*, *B. bronchiseptica*, and *B. parapertussis* (34). A reanalysis of S17, in which the identity threshold was lowered, revealed that a 27 nt region starting at the 5’-end of the S17S (highlighted in yellow in Fig. 2A and B) is 100% conserved more broadly, in species throughout the *Burkholderiales* order and even within other species of *Betaproteobacteria* (Fig. 3). The conservation of a predicted factor-independent terminator at the 3’-end suggests that these putative S17S homologs are generated similarly to S17S in *B. pertussis*. These results suggest a possible conserved function for this particular region of S17S among hundreds of bacterial species.

**FIG 3.**
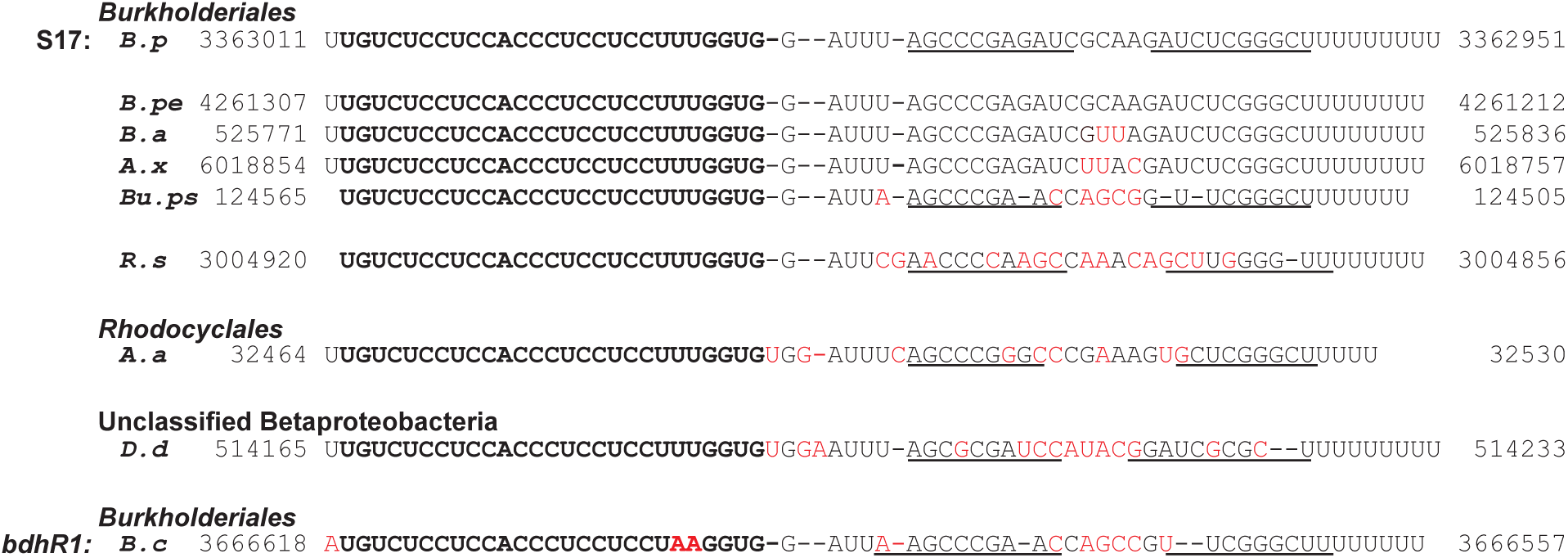
The 5’-end of S17S is highly conserved among a subset of Betaproteobacteria. The top line shows the RNA sequence of S17S in *B. pertussis* (*B.p*) from nucleotides +124 to +190 relative to the +1 TSS (positions 3363011 to 3362951 in *B. pertussis* Tohama I NC_002929.2) (See Fig. 2). Underneath are shown sequences with the genomic positions indicated from the following species in the *Burkholderiales* order: *Bordetella petrii* strain DSM 12804 (*B.pe*, Sequence ID AM902716.1), *Bordetella avium* 197N (*B.a*, Sequence ID AM167904.1), *Achromobacter xylosoxidans* A8 (*A.x*, Sequence ID CP002287.1), *Burkholderia pseudomallei* 1026b (*Bu.ps*, Sequence ID CP004379.1), and *Ralstonia solanacearum* T12 (*R.s.*, Sequence ID CP022774.1); from the following species in the *Rhodocyclales* order: *Aromatoleum aromaticum* EbN1 (*A.a*, Sequence ID CR555306.1); from the unclassified *Betaproteobacterium Candidatus desulfobacillus denitrificans* (*D.d*, Sequence ID AP021857.1); and from *Burkholderia cenocepacia* J2315 (*B.c*, Sequence ID AM747720.1 chromosome 1, NC_011001.1 chromosome 2, and NC_011002.1 chromosome 3) in the *Burkholderiales* order. The identical sequences found in the top 8 species are shown in bold black with the mismatches in *B.c* shown in bold red. Conserved and non-conserved nucleotides downstream of this region are indicated in regular black and red, respectively. The position of the stem-loop at the 3’-end is indicated by the lines.

The S17 sequence also encodes a putative small protein of 32 residues, but the 5’-end of S17S lies within the open reading frame (ORF). Consequently, if synthesized, the complete peptide could only be generated from the S17L RNA.

We previously found that the level of S17, particularly S17S, is significantly higher in the absence of MgSO_4_, suggesting that its expression is enhanced by the presence of BvgA∼P (34). In addition, S17S coprecipitated with a FLAG-tagged *B. pertussis* Hfq, indicating a direct interaction between S17 and this RNA chaperone (34). Northern analyses using RNA isolated from isogenic *B. pertussis* WT, *ΔbvgAS*, or *Δhfq* strains showed a significant decrease in the level of the sRNA in the mutant strains either in the presence or absence of MgSO_4_, affirming the enhancement of S17 by BvgA∼P and Hfq (Fig. 1B).

### S17L is generated by transcription from a σ^A^-dependent promoter

Transcription from the predicted strong σ^A^-dependent promoter P_S17_ would generate the S17L species. To ask whether S17L is indeed an initiated mRNA, we treated total RNA extracts from WT *B. pertussis* grown with and without MgSO_4_ with Terminator™ 5ߣ-Phosphate-Dependent Exonuclease (TEX), which degrades RNAs containing a 5’-monophosphate. Thus, processed RNAs are susceptible to TEX, while transcription-initiated mRNAs, which have a 5’-triphosphate, are resistant. After incubating the RNA with and without TEX, both S17L and S17S were observed in the absence of TEX, while the S17S transcript was eliminated in the presence of TEX (Fig. 4A). As expected, the level of the control 5S transcript was not affected by TEX (Fig. 4A). Taken together, these results suggest that S17L arises from P_S17_ transcription while processing of S17L generates S17S.

**FIG 4.**
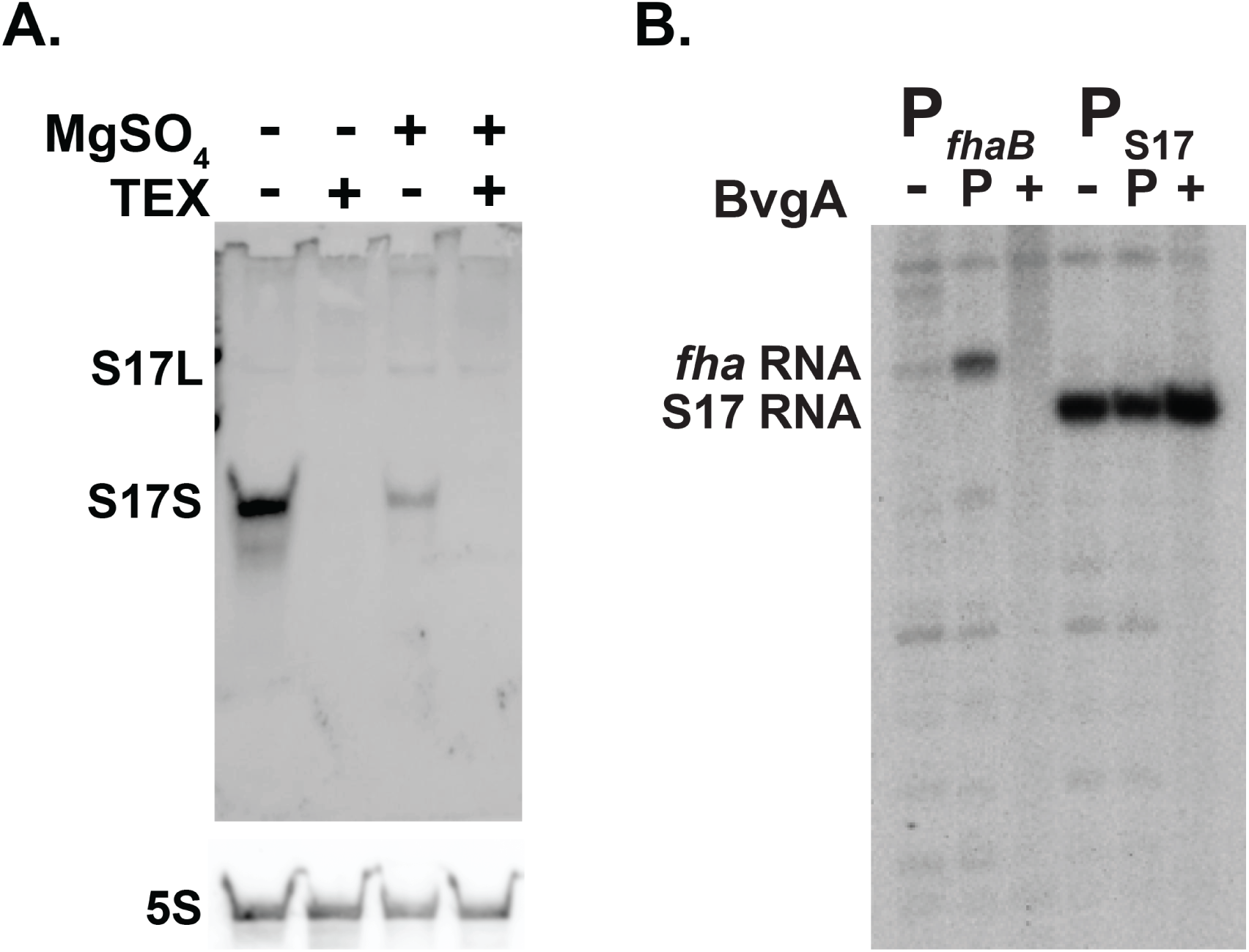
S17S is a processed RNA, while S17L is generated as a transcript by *B. pertussis* σ^A^-RNAP. (A) Representative Northern blot (one of 2 biological replicates) showing the S17 products with and without treatment of RNA isolated from WT *B. pertussis* grown with and without MgSO_4_ and with and without treatment with 5’ monophosphate dependent terminator exonuclease (TEX); underneath a rehybridization of the gel with a probe for 5S RNA is shown. (B) Representative denaturing, polyacrylamide gel (one of 2) showing the *in vitro* transcription products obtained using *B. pertussis* σ^A^-RNAP and the indicated supercoiled DNA harboring the BvgA∼P-dependent promoters P*_fhaB_* or P_S17_. Transcriptions were performed in the absence of BvgA (-) or in the presence of BvgA (+) or BvgA∼P (P).

To ask whether the P_S17_ transcript is generated by σ^A^-RNAP, we performed *in vitro* transcription reactions using purified *B. pertussis* σ^A^-RNAP and a supercoiled plasmid harboring P_S17_ from −175 to +86 relative to the transcription start site (TSS). In this construct, transcription from P_S17_ will result in an RNA of ∼365 nt after termination at the downstream T7 terminator present in the vector. As a control, we used the BvgA∼P-dependent promoter P*_fhaB_*, which yields a transcript of ∼ 400 nt. As seen in Fig. 4B, P_S17_ generated an RNA consistent with the expected species.

Given that S17 has been classified as a Bvg^+^ mode gene [(34); Fig. 1B], we investigated whether the presence of BvgA∼P increased the level of P_S17_ RNA *in vitro*. While transcription from P*_fhaB_* was strongly dependent on the addition of BvgA∼P as expected, transcription from P_S17_ was unchanged by the presence of either non-phosphorylated BvgA or BvgA∼P (Fig. 4B). Thus, P_S17_ is a σ^A^-dependent promoter that does not require BvgA for activity. We postulate that P_S17_ is repressed *in vivo* in the Bvg^-^ mode and that the increase in S17 abundance in the Bvg^+^ mode arises from derepression of P_S17_ transcription.

### Processing of the unstable 5’-triphosphated S17L by RNase E generates stable S17S from the unstable S17L

To investigate the stability of S17 *in vivo*, we observed the amounts of S17L and S17S present after treating exponentially growing *B. pertussis* Tohama I BP536 cultures with rifampicin, which inhibits transcription initiation. Cultures were grown without MgSO_4_, a condition in which S17S is abundantly expressed (Fig. 1B). While S17L disappeared quickly (half-life of < 2 min), S17S was significantly more stable; ∼ 50% remained after 20 min (Fig. S1A). As expected, the control sRNA SsrA was stable (Fig. S1B). These results suggested that the abundant S17S species arises from processing of S17L.

Previous work has implicated both RNase III and RNase E as enzymes that are responsible for much of the RNA processing in Gram-negative bacteria (38, 39). As the *B. pertussis* RNase III and RNase E proteins are homologous with those from *E. coli*, we used a set of *E. coli* strains to investigate whether RNase III and/or RNase E were involved in S17 processing: WT, Δ*rnc* (lacking RNase III), *rne*^ts^ (containing a temperature sensitive RNase E), and Δ*rnc,rne*^ts^. We cloned the S17 gene in a plasmid downstream of the inducible *lac* promoter and then induced expression of the sRNA in each *E. coli* strain before and after a temperature switch to inhibit RNase E activity. The steady-state levels of S17L or S17S were then obtained from Northern analyses and normalized with those obtained using the WT (Fig. 5). It should be noted that the level of S17L relative to S17S was higher in these strains compared to what we observed in *B. pertussis* (Fig. 1B).

**FIG 5.**
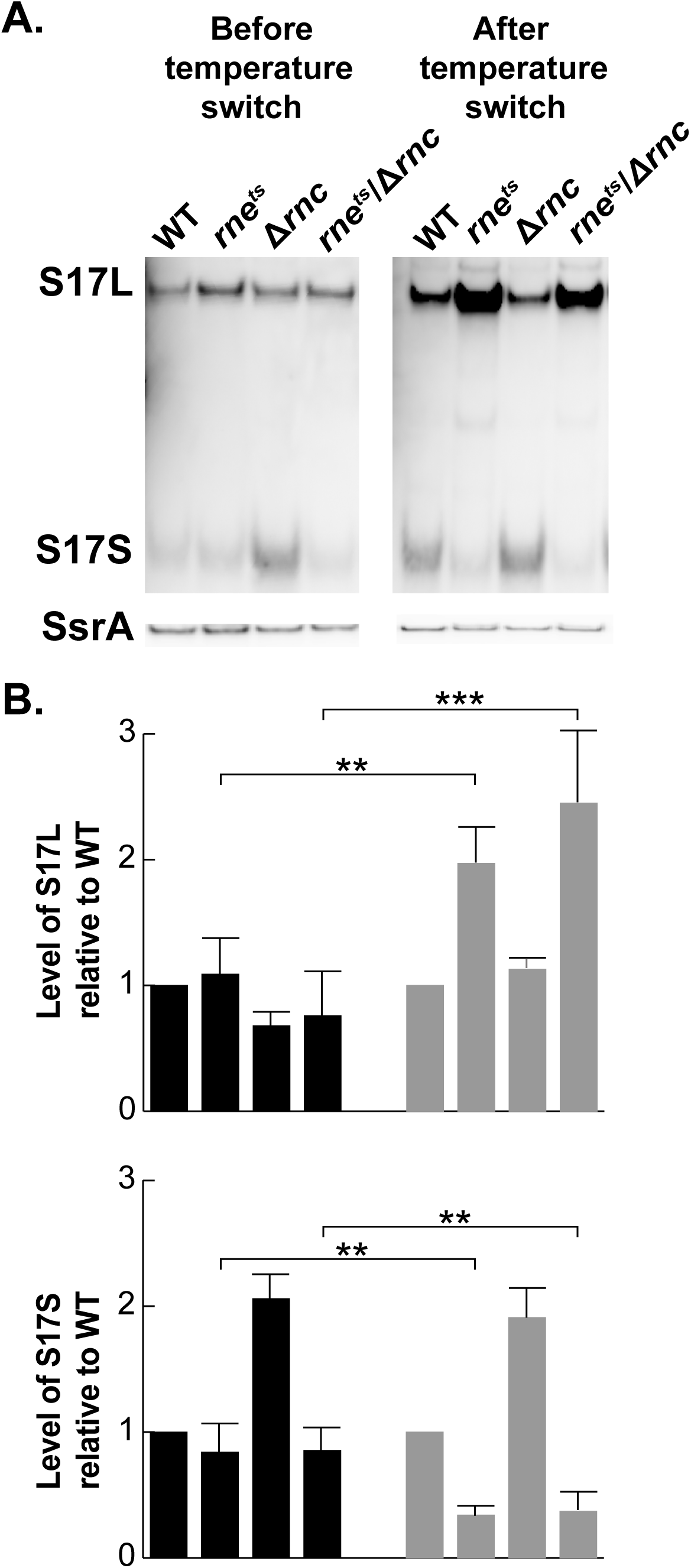
S17L is processed by RNase E to generate S17S. Representative Northern blots (one of three biological replicates) show S17L and S17S, before and after a 15 min temperature switch from 30° C to 42.5° C to inactivate the temperature sensitive RNase E mutant protein present in the indicated *rne^ts^*strains (38); S17 was generated from pGFK3038, which contains S17 downstream of P*_lac_*. A rehybridization of the gel with a probe for the sRNA SsrA is shown below. Plots show the quantification of the levels of S17L and S17S in the various RNase mutants relative to WT from the Northern blot analyses. Error bars indicate the standard deviation from the biological triplicates. Student t-test was performed to obtain statistical significance. Significant differences with p-values of <0.01 (**) and <0.001 (***) are indicated.

Before the temperature shift, the levels of S17L were similar in all the strains, while the steady state level of S17S was similar in all the strains except for Δ*rnc* where it was ∼2-fold higher, suggesting that the stability of the processed S17S may be affected by RNase III. In contrast, 15 min after the temperature switch, the level of S17L increased ∼2-fold while the level of S17S decreased ∼2-fold in both the *rne*^ts^ strain and the Δ*rnc,rne*^ts^ double strain. These results are consistent with the hypothesis that S17S arises through processing of S17L by RNase E. Furthermore, the site of S17 cleavage (5’GU↓UGU3’) is similar to the ideal RNase E cleavage site (5’RN↓WUU3’) (40).

### Identification of possible *B. pertussis* genes targeted by S17

Taken together, our analyses argued that S17S is an RNase E processed sRNA, which interacts with Hfq and whose level significantly increases in the presence of BvgA∼P. Consequently, it seemed likely that an interaction of S17S with specific mRNAs, mediated by Hfq, might result in the regulation of specific genes in the Bvg^+^ mode.

To investigate whether particular genes might be affected by S17, we generated an isogenic ΔS17 strain from the WT strain, *B. pertussis* Tohama I BP536, and isolated RNA from the WT and ΔS17 strains grown under nonmodulating conditions (no MgSO_4_; Bvg^+^ mode). This is the mode in which the level of S17S is enhanced. We then performed RNA-seq analyses using a total of 7 biological replicates, with datasets obtained on different days, from 2 (dataset 1), 2 (dataset 2), or 3 (dataset 3) independent RNA isolations. (Transcriptomic data are available in the NCBI database GEO#: GSE277013 and Table S1.) In each dataset, a log2fold change (FC) > 0.7 in gene expression with an adjusted *p*-value < 0.05 after an Empirical edgeR statistical test was scored as significant.

As many sRNAs are involved in translational repression (2), we focused on genes whose levels increased in the ΔS17 strain relative to WT. We looked for genes that had a log2FC > 0.7 in at least 1 of the RNA-seq datasets, and as an independent analysis, we performed RT-qPCR for 25 genes to test the RNA-seq data. Using these criteria, we identified 91 possible S17 potential targets (Table S2; shown as green dots in Fig. S2).

Investigation of the identified genes using a published transcriptome analysis for *B. pertussis* (33) indicated that some of the identified genes were present within operons, while others represented single transcription units (Table S2). Since many sRNAs are known to repress translation by annealing to a sequence near the start of a gene, we searched either the 5’UTRs identified from the TSSs or the intergenic regions upstream of genes’ translation start sites to ask whether there were possible annealing sites for S17S. This analysis indicated that 70 genes (77%) of those identified by the RNA-seq/RT-qPCR analyses contained significant matches to the single-stranded (ss) regions of S17S: the 5’-end (designated as Site 1), the first predicted loop (Site 2), or Site 1 plus the second predicted loop (Site 3) (Table S2 and Fig. 6A). As noted above, the sequence from positions 2-28 of S17S, which includes Sites 1 and 2, is totally conserved in species throughout the Betaproteobacteria class of bacteria (Fig. 3). In nearly all of the genes, annealing to either Site 1 or Site 2 should disrupt ribosome binding. The identified genes encode 32 transporter-related genes/operons, 3 transcriptional regulators [*BP0441* (MerR family), *BP2158* (sigma 54-dependent activator), and BP3670 (TetR/AcrR family], 4 potential metal binding proteins [(*BP0441* (MerR family regulator), *BP0442* (MBL fold metallo-hydrolase), *BP1555* (zinc metallopeptidase), and *BP2864* (zinc-binding dehydrogenase), nearly all the genes involved in phenylacetic acid catabolism (*paa* genes), and various reductases, hydrogenases, and hydrolases (Table S2).

**FIG 6.**
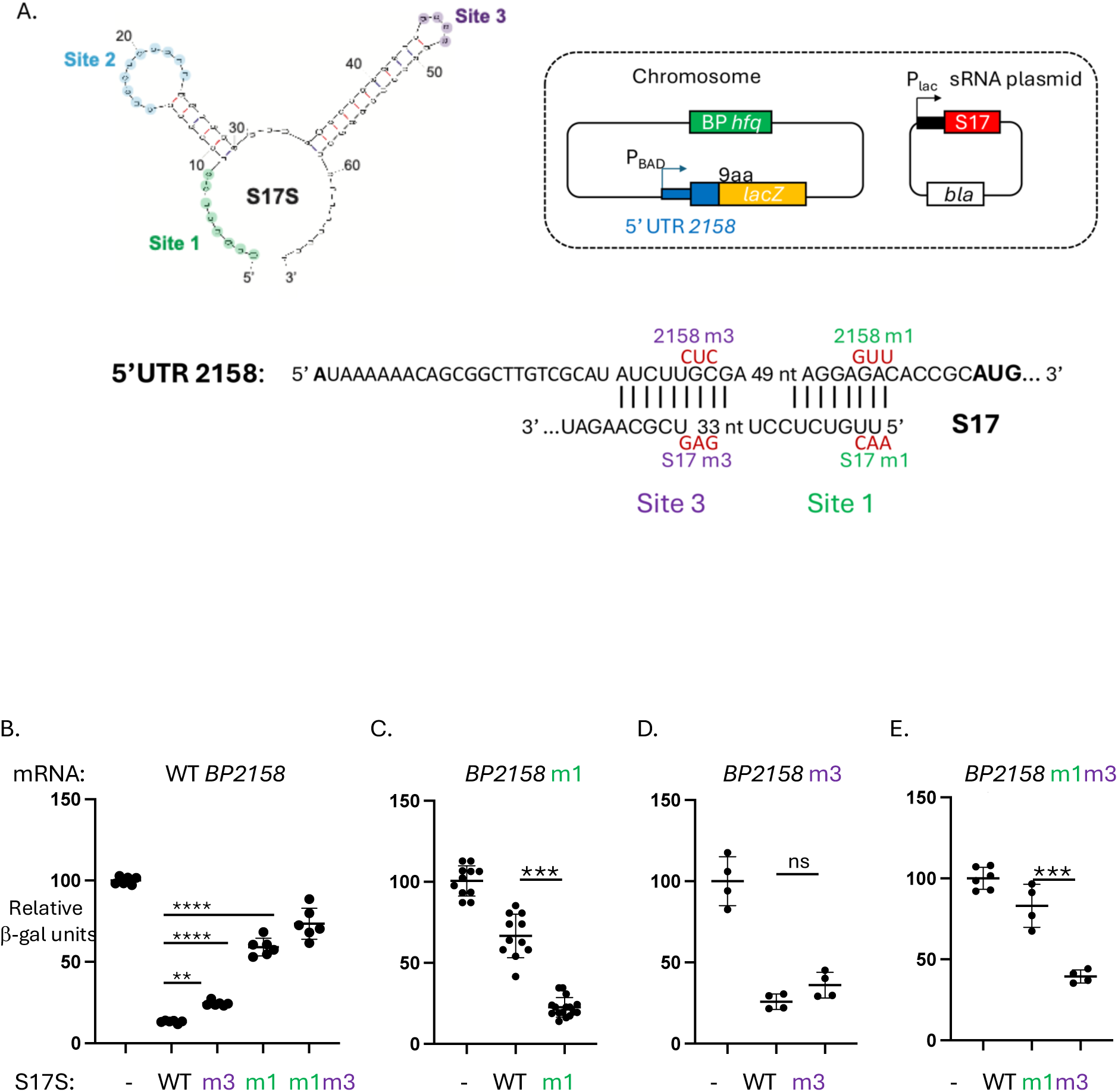
S17S post-transcriptionally represses *BP2158.* (A) Translational reporter assay. Top left, mfold predicted structure of S17S with the single-stranded regions of Sites 1, 2, and 3, shown in green, blue, and purple, respectively. Top right, schematic of translational reporter assay for *BP2158*. In the *E. coli* chromosome, *E. coli hfq* was replaced with FLAG-tagged *B. pertussis* (*Bp*) *hfq* (green) and a *lacZ* reporter gene (yellow) was fused to the 5’UTR of *BP2158* plus its first 9 codons (blue). The *BP2158-lacZ* fusion is transcribed from the arabinose inducible promoter, P_BAD_. An ampicillin-resistant (*bla*) multicopy plasmid contains either S17S or S17L (red). The vector without the S17 insert was used as the control. Bottom, Sequence of the *BP2158* 5’UTR in black with the AUG translation start in bold. The sequences that can potentially anneal to S17 Sites 1 and 3 are indicated. Above the BP2158 sequence and below the S17 sequence are shown in red the mutations in either BP2158 5’UTR or S17, respectively, that disrupt annealing to Sites 1 and 3. (B-E) Results of β-galactosidase (β-gal) assays showing the relative β-gal (Miller) units using the indicated S17S plasmids in the strains WT BP2158 (B), BP2158m1 (C), BP2158m3 (D), and BP2158m1m3 (E), relative to the plasmid without the S17S insert (-). Means and standard deviations are shown; in some cases, the points are too close together to be individually distinguishable. Results of one-way ANOVA comparison tests for various datasets are indicated: NS, not significant; **, p-value < .01; ***, p-value < .001; ****, p-value < .0001.

Given that S17 is an Hfq-enhanced sRNA [(34) Fig. 1B], it seemed likely that the ability of S17 to interact with target genes is dependent on the formation of an mRNA/Hfq/S17 complex. To investigate this, we asked whether the identified up-regulated genes overlapped with genes previously found to be up-regulated by a deletion of *hfq* in *B. pertussis* Tohama I (12). We found that 50 genes (55%) were significantly affected in the Δ*hfq vs.* WT microarray datasets representing exponential phase growth, stationary phase growth, day zero of mouse infection, and/or 12 days post-infection (Table S2). This correlation is consistent with their regulation by an sRNA that involves Hfq.

### Overexpression of S17 directly represses *BP2158*, a predicted σ^54^ transcriptional regulator

In the ΔS17 strain the expression of *BP2158*, one of two predicted σ^54^ transcriptional regulators in *B. pertussis*, increased relative to the WT strain in two out of the three RNA-seq datasets, and RT-qPCR analysis confirmed this increase (Table S2). *BP2158* encodes a predicted FIS family regulator of σ^54^-RNAP and is present as a single ORF with an assigned TSS 92 nt upstream of the AUG start (33). The *BP2158* 5’UTR contains extensive complementarity with S17S, having an 8/8 match with Site 1 and a 9/9 match with Site 3 (Fig. 6A).

To determine whether S17 could post-transcriptionally repress expression of *BP2158*, we modified an *E. coli* system, which had previously been employed to investigate the effect of sRNA overexpression on translation *in vivo* [(41)(Fig. 6A)]. In this system, we replaced the *E. coli hfq* gene with a FLAG-tagged *B. pertussis hfq* construct, since we had previously demonstrated that Hfq^Bp^-FLAG is active in *E. coli* (34). We then constructed a strain in which the arabinose-inducible promoter P*_BAD_* was positioned upstream of the 5’UTR/start of *BP2158* (through the first 9 codons) fused in frame to *E. coli lacZ*. The resulting strain was transformed with the vector pBR-plac, which has the IPTG-inducible promoter P*_lac_* but no insert, or with pS17L or pS17S, in which the sequence of S17L or S17S was inserted into pBR-plac downstream of P*_lac_*, respectively (Fig. 6A).

We performed β-galactosidase assays from extracts obtained in the presence of 0.1 mM IPTG to induce expression of the fusion reporter genes and 0.02% arabinose to induce expression of S17. We found that overexpression of S17S or S17L repressed translation of *BP2158* by ∼ 7.5- fold or ∼4-fold, respectively (Fig. 6B, Fig. S3).

To investigate whether annealing between S17 Site 1 and/or Site 3 was indeed important for repression, we constructed strains with 3 nt changes within the *BP2158* 5’UTR, designated as *BP2158*m1, *BP2158*m3, and *BP2158*m1m3. These mutations would substantially disrupt the ability of S17 to anneal with the *BP2158* 5’UTR using Sites 1, 3, or 1 and 3, respectively (Fig. 6A). We also constructed plasmids, containing the complementary changes in S17S, designated as pS17Sm1, pS17Sm3, and pS17Sm1m3 (Fig. 6A).

Disruption of the annealing at Site 1 using either *BP2158*m1 or S17Sm1 significantly disrupted repression (Fig. 6B, C), while the presence of both *BP2158*m1 and S17Sm1 restored repression (Fig. 6B). Although disruption of annealing at Site 3 alone had only a slight effect (Fig. 6B, 6D), the results with having both Sites disturbed (Fig. 6B) or restored (Fig. 6E) suggested that Site 3 may enhance the effect of Site 1. Similar results were observed when using WT and mutant pS17L plasmids with the WT *BP2158* strain, except that the level of repression by S17L was about 2-fold less than that of S17S (Fig. S3). From these results we conclude that S17 directly represses BP2158 post-transcriptionally.

### Effect of S17S on other candidate genes

Using the same modified *E. coli* system, we also investigated possible repression by S17 of other candidates: *BP2642A*, a gene at the 5’-end of an ABC transporter operon with complementarity to S17 Sites 1 and 3; *BP1447* (*mmsB*, a 3-hydroxyisobutyrate dehydrogenase, complementarity to Site 1); *BP0442* (a metallo-hydrolase, complementarity to Sites 1 and 3); and *BP1445* (an acyl-CoA dehydrogenase family protein, complementarity to Site 1) (Table S2). In the case of *BP2642A*, several downstream genes (*BP2642*-*BP2638*) were also expressed significantly higher in the S17 deletion strain, suggesting that S17 might regulate the entire operon.

Although we were unable to detect an effect of S17S or S17L overexpression on the expression of *BP0442* or *BP1445*, overexpression of S17S reduced the level of the *BP2642A*-*lacZ* and the *mmsB*-*lacZ* fusion proteins by ∼2-fold (Fig. S4), suggesting that S17S also regulates the *BP2462A*-*BP2638* region and *mmsB*. In contrast, overexpression of S17L had no effect on *BP2642A* and *mmsB* (Fig. S4).

### Possible conservation of S17S regulation

Our analyses described above revealed that various Betaproteobacteria species have a potential sRNA with 100% identity to S17S positions 2-28, which includes Sites 1 and 2 (Fig. 3). To investigate the possible conservation of S17 regulation, we asked whether 11 S17 targets, like S17, were also conserved in a range of Betaproteobacteria: *Bordetella bronchiseptica* (*B.b*) and *Achromobacter xylosoxidans* (*A.x*) (*Alcaligenaceae* family, *Burkholderiales* order), *Ralstonia solanacearum* (*R.s, Burkholderiaceae* family*, Burkholderiales* order), and *Aromatoleum aromaticum (A.a, Rhodocyclaceae* family, *Rhodocycales* order). We found potential S17S regulation for 4 out of 10 of the selected targets in all the tested species, while 4 out of 10 targets extended within the *Alcaligenaceae* family of the *Burkholderiales* order (Tables 1 and S2). These results support the idea that regulation of multiple genes by S17S is a conserved process.

### S17 has significant homology to ‘toxic sRNAs’ present in *Burkholderia cenocepacia*

In addition to the 100% conservation of the 5’-end of S17S (Sites 1 and 2) to sRNAs found throughout a range of betaproteobacteria, another family of sRNAs identified in *Burkholderia* species also have excellent matches to the 5’-end of S17 (42–44) (Fig. 3). One particular sRNA, designated as *bdhR1* (Rfam ID RF02278), has only a 2 nt mismatch with S17 in this region. This family has been referred to as ‘toxic sRNAs’ since their overexpression significantly suppresses *E. coli* or *B. cenocepacia* growth in rich media. However, for *E. coli* BL21 grown in LB medium (Fig. S5), overexpression of S17S did not impair exponential growth and only modestly impaired growth upon entering stationary phase, indicating that under these conditions S17 is not similarly toxic.

*bdhR1* is more highly expressed during starvation, in growth in glucose medium, and in biofilm or stationary phase compared to planktonic growth [reviewed in (45)]. Bioinformatic analysis revealed ∼100 genes as potential *bdhR1* targets, including a Fis family regulator of σ^54^- RNAP (Bcam2365) (Table S2). Our visual analysis revealed two other Fis family regulators of σ^54^-RNAP (Bcam2211 and Bcam2715). The 5’UTRs of all three regulator genes have excellent matches to the Site 1 target sequences similar to that of *BP2158*. Taken together, this suggests some overlapping roles for S17 and *bdhR1*.

## DISCUSSION

Regulation of gene expression in bacteria by sRNAs serves to fine-tune the level of gene expression by enhancing/interfering with nuclease cleavage of mRNA or by improving/interfering with ribosome binding (1–3). In pathogenic bacteria, sRNA regulation has been tied to virulence through its modulation of the level of quorum sensing, Type III secretion systems (T3SS), iron transport, and biofilm formation; *Δhfq* strains are often impaired in these functions, suggesting that the sRNA chaperone Hfq is also involved in this regulation (2, 4–8, 11, 12, 46). However, despite the documented importance of sRNAs and Hfq in bacterial virulence overall, their relevance to *B. pertussis* has been less explored. Previous work has suggested that sRNAs may play an important role for this pathogen. Hundreds of sRNAs have been predicted from transcriptomic analyses (34, 36, 47), and deletion of *hfq* from the *B. pertussis* genome attenuates the expression of various virulence genes, including those within the T3SS (12, 32). In addition, two sRNAs in *B. pertussis* have previously been characterized: RgtA and Bpr4. RgtA is a Bvg^-^ mode sRNA, which is dependent on the RisA response regulator and regulates the expression of *BP3831*, a gene involved in glutamate metabolism (35). Bpr4 is a Bvg^+^ mode, activating sRNA that up-regulates *fha* (adhesin filamentous hemagglutinin) during tracheal colonization (37).

Here we have characterized S17, the first BvgA^+^ mode, Hfq-dependent repressing sRNA identified for *B. pertussis*. The long form of S17 initiates from a σ^A^-dependent promoter and terminates at a factor-independent transcription terminator, while a much more abundant short form arises from RNase E-dependent processing of S17L. Our work indicates that S17 directly down-regulates the σ^54^ transcriptional activator *BP2158* and suggests that multiple additional genes are targeted, including transporters and metal-binding proteins. The 5’-end of S17S is 100% conserved in a range of Betaproteobacteria orders (Fig. 3) and potential S17 targets are also conserved (Table 1), suggesting an evolutionarily conserved function for this sRNA among hundreds of bacterial species. In addition, the similarity between S17 and the characterized *B. cenocepacia* sRNA *bdhR1* suggests that like *bdhR1*, it may be involved in regulating genes related to survival and metabolic changes. Certainly, the repression of genes in the Bvg^+^ mode would be consistent with its attenuating genes that are needed in the Bvg-mode, which involves survival outside of the respiratory tract (16, 26). However, the lifestyle and genomic capacity of *B. pertussis* and *B. cenocepacia* differ greatly. While *B. pertussis* is an obligate human pathogen with a relatively small genome, *B. cenocepacia*, with a much larger genome size, is found in a wide range of natural environments and acts as an opportunistic pathogen in humans, particularly among cystic fibrosis patients (48). Thus, there are likely to be many differences between the functions of S17 and *bdhR1*.

**Table 1.** Conservation of S17 targets within Betaproteobacteria.

| Gene<sup>a</sup> | Function<sup>b</sup> | S17 | Present<sup>d</sup> in |  |  |  |
| --- | --- | --- | --- | --- | --- | --- |
|  |  | Binding Site<sup>c</sup> | *B.b.* | *A.z.* | *R.s.* | *A.a.* |
| *BP0441* | MerR family regulator | Site 2 | + | + | - | - |
| *BP0596* | transporter permease | Site 2 | + | + | none<sup>e</sup> | - |
| *BP1447* | MmsB dehydrogenase | Site 1 | + | - | - | none<sup>e</sup> |
| *BP1487* | periplasmic solute binder | Site 1 | + | - | none<sup>e</sup> | + |
| *BP2158* | σ<sup>54</sup> regulator | Site 1 | + | + | + | + |
| *BP2271* | extracellular solute binder | Site 1 | + | - | - | none<sup>e</sup> |
| *BP2418* | transporter binder | Site 1 | + | + | + | + |
| *BP2642A*<sup>f</sup> | transport ATP binder | Site 1 | + | + | + | +<sup>g</sup> |
| *BP2678* | PaaN phenylacetic acid degradation | Site 1 | + | + | none<sup>e</sup> | +<sup>g</sup> |
| *BP2811* | fatty acid ligase | Site 1 | + | + | + | + |
| *BP3501* | transporter binder | Site 1 | + | + | - | none<sup>e</sup> |

<sup>a</sup>gene name in *B. pertussis* Tohama I; <sup>b</sup>predicted or identified function; <sup>c</sup>potential annealing site within S17, see Table S2 for sequences; <sup>d</sup>*B.b.*, *A.z.*, *R.s.*, and *A.a.*: *Bordetella bronchiseptica* 253, *Achrombacter xylosoxidans* A8, *Ralstonia solanacearum* T12, and *Aromatoleum aromaticum* EbN1, respectively; <sup>e</sup>gene homolog was not found; <sup>f</sup>while *BP2642A* is a pseudogene in *B. pertussis*, the homologs are actual genes and in each case the homolog, like *BP2642A*, is at the beginning of an operon; <sup>g</sup>binding is to Site 2 rather than Site 1.

The finding that S17 directly represses a σ^54^ transcriptional regulator is particularly intriguing since the role of this alternative σ factor in *B. pertussis* has not yet been explored. Sigma factors encode the specificity subunits for bacterial RNAP, recognizing and binding specific DNA sequences in promoter regions to initiate transcription (49, 50). All bacteria have a major ‘housekeeping’ sigma factor, σ^A^, such as σ^70^ in *E. coli*, which is required for growth. However, most bacteria, including *B. pertussis*, also have alternative σ factors, including σ^54^, which are used for various specific conditions and/or stressors.

σ^54^ is unique in that it absolutely requires a transcriptional activator ATPase for activity (51, 52). Consequently, each activator is responsible for gene(s) that are under the control of a particular σ^54^ regulon. In *B. pertussis* there are 2 such σ^54^ factors, encoded by *BP2158* and *BP2004*. Our previous RNA-seq analyses have indicated that *BP2004* is not affected by *bvgAS*, but the level of *BP2158* decreases approximately 3-fold in the Bvg^+^ mode (25). This is consistent with our finding that S17 represses *BP2158* in the Bvg^+^ mode. Furthermore, a visual analysis of the sequence upstream of *BP2158* indicates the presence of a nearly perfect σ^54^ promoter sequence (Table S2), located ∼75 bp upstream of the ATG start. If active, this promoter would require a σ^54^ activator, perhaps BP2158 itself, in addition to σ^54^-RNAP.

Taken together, our results suggest a complicated pattern for the regulation of a set of BP2158/σ^54^-dependent gene(s) (Fig. 7). In the Bvg^+^ (virulence) mode, the level of S17S is high because transcription from the σ^A^-dependent promoter P_S17_ yields S17L, which is then processed by RNase E to produce an abundant level of the stable, short S17S form. The resulting high level of S17S, in turn, lowers the expression of the BP2158/σ^54^-dependent gene(s). However, in the Bvg^-^ mode, the mode associated with transmission and persistence, the level of S17 is low, resulting in the expression of the BP2158/σ^54^-dependent regulon. Since P_S17_ does not need an activator for activity, we speculate there must be a repressor of P_S17_ in the Bvg^-^ mode, which is itself repressed in the Bvg^+^ mode. Finally, transcription of *BP2158* itself appears to arise from a σ^54^-RNAP promoter, suggesting that BP2158 may activate its own transcription.

**FIG 7.**
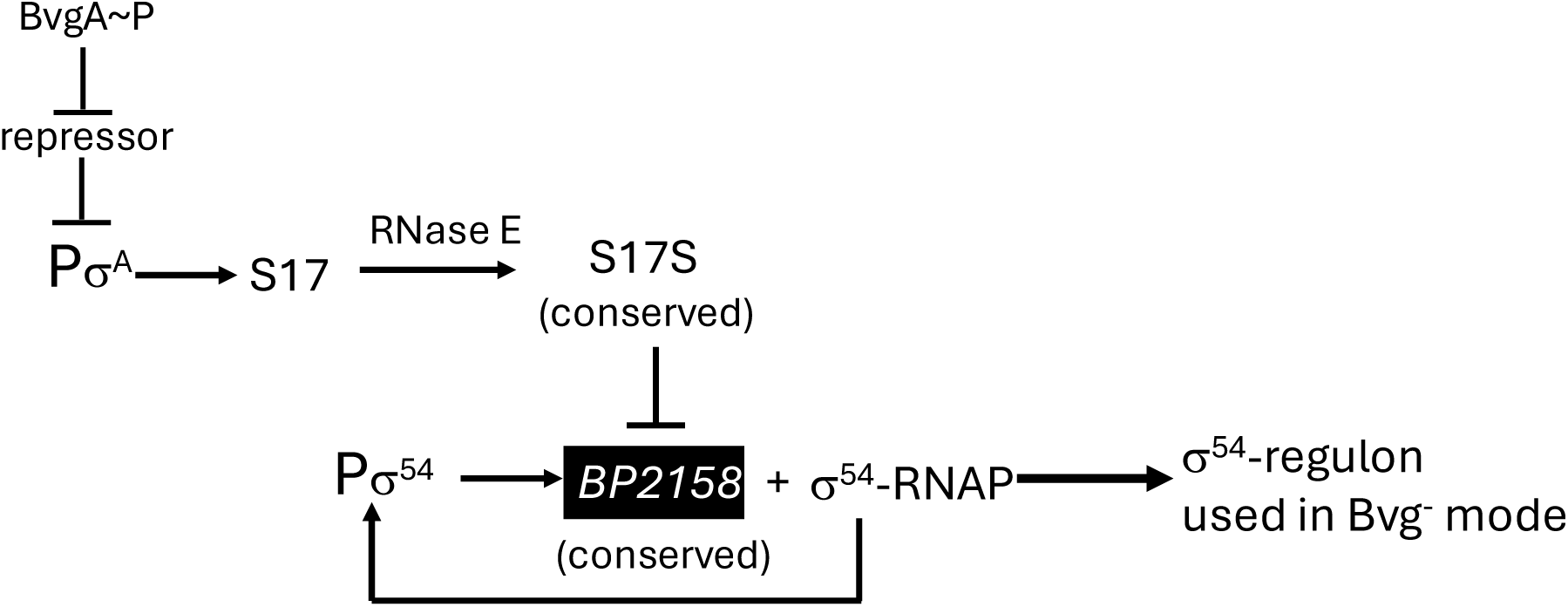
Proposed model for the regulation of a set of Bvg^-^ genes by S17S and the σ^54^ transcriptional activator BP2158 together with σ^54^-RNAP. The sRNA S17 is expressed by the σ^A^-dependent promoter P_S17_, whose repression is relieved in the Bvg^+^ mode when BvgA∼P is present. This leads to expression of S17, which is then processed by RNase E to generate an abundant level of S17S, which then represses *BP2158* post-transcriptionally. However, in the Bvg^-^ mode when BvgA∼P is not present, P_S17_ is repressed, lowering the level of S17, leading to an increase in the level of *BP2158*. An excellent match to the consensus promoter sequence upstream of *BP2158* suggests that it is expressed by σ^54^-RNAP.

The role of σ^54^ and the gene(s) within the *BP2158*/σ^54^ regulon in *B. pertussis* are not yet known. σ^54^ was originally designated as a ‘nitrogen’ sigma, needed for nitrogen metabolism, but now it is known to be involved in several processes in bacteria, including motility, biofilm formation, and membrane stress (53–55). Very recent work has shown that a knockout of *rpoN* encoding σ^54^ in *B. bronchiseptica* RB50, which shares 99% identity with *B. pertussis rpoN*, decreases motility, biofilm formation, and the fitness of *B. bronchiseptica* to colonize rat trachea, indicating the importance of σ^54^ in *B. bronchiseptica* infection (56). *B. bronchiseptica* contains several σ^54^ transcription regulators, but one is 99% identical to *BP2158*. Thus, we speculate that a *BP2158*/ σ^54^-RNAP regulon present in both *B. pertussis* and *B. bronchiseptica* may have similar roles. Further work will be needed to identify this regulon and how S17 contributes to this and other *B. pertussis* pathways.

## MATERIALS AND METHODS

### Bacterial strains and cell culture

*B. pertussis* Tohama I BP536 (57) and the derivatives Δ*bvgAS* (25) and Δ*hfq* (34) have been described. The ΔS17 derivative was constructed as follows. Plasmid pQC2473 was created by inserting a 926 bp synthetic fragment into the allelic exchange vector pSS4894 (23). This fragment was comprised of two 460 bp DNA segments flanking the region to be deleted (bp 3363169 – 3362945 of the *B. pertussis* Tohama I genome, GenBank NC_002929.2) with an *EcoR*I site at the junction of the two fragments. Subsequently, an 866 bp synthetic fragment encoding spectinomycin resistance was cloned as an *EcoR*I fragment into the *EcoR*I site of pQC2473 to create plasmid pQC2474. This plasmid was transformed into the *E. coli* donor strain RHO3 (58), with selection for gentamicin resistant colonies on LB agar supplemented with 200 µg/mL 2,6- diaminopimelic acid (DAP) and 10 µg/mL gentamicin. Allelic exchange was performed using the Tohama I derivative BP536 as a recipient and RHO3/pQC2474 as a donor in a single mating step with selection for spectinomycin resistance (50 µg/mL) to create strain QC5169 (BP536, ΔS17::spcR). The marked S17 deletion in QC5169 was replaced by allelic exchange with RHO3/pQC2473 as donor to create the strain QC5170 (BP536, ΔS17), using a two-step allelic exchange procedure as previously described (23).

*B. pertussis* strains were grown at 37° C in Bordet Gengou (BG) media (59) or BG agar containing streptomycin (50 μg/mL) in the absence or presence of 50 mM MgSO_4_ as described (Chen *et al.*, 2010).

*E. coli* DH5σ (60) and *XL1-Blue* (Agilent, Santa Clara, CA, USA) were used for standard cloning procedures, and *E. coli TOP10* (Agilent) was used for TOPO-cloning in the 5’-RACE analysis. BL21 competent cells were obtained from New England BioLabs (NEB, Ipswich, MA, USA).

*E. coli* chromosomal mutant strains used for *lacZ* reporter assays were constructed from *PM1805* (*MG1655 mal::lacI^Q^ ΔaraBAD araC^+^ lacI’::PBAD-cat-sacB-lacZ, miniλ-tetR*) (41) by phage P1-mediated transduction using KM3033 (34), which contains a FLAG-tagged *B. pertussis hfq* rather than *E. coli hfq*, as the donor strain. From this construction, a set of strains were generated as described using gBlocks (Integrated DNA Technologies, Newark, NJ; See Table S3 for sequences) (61), resulting in a chromosomal gene fusion of the first nine codons of the *B. pertussis* gene to the 10^th^ codon of the *lacZ* gene; the fusion gene was located downstream of the arabinose-inducible P_BAD_ promoter and the 5’UTR represented the *B. pertussis* sequence either from the assigned TSS (47) or from the intergenic region upstream of the *B. pertussis* gene. The indicated mutant strains were generated similarly using gBlocks in which the predicted annealing sites for S17 were changed as indicated (Fig. 6A). Recombinants were selected on M63 media (62) containing 5% sucrose, 0.2% glycerol. The DNA regions containing the *B. pertussis* inserts were sequenced by Psomagen, Rockville, MD.

For RNase assays, the following *E. coli* strains were used: WT EM1055 [DJ480: MG1655 Δ*lac* X174 (63)] and the derivatives of EM1055: EM1277 [*rne*-*3071^ts^ zce-726*::Tn*10* (38)], which contains a temperature-sensitive mutant of *rne*, encoding RNase E; EM1321 [EM1055 *rnc*- *14*::Tn*10* (38)], which contains a deletion of *rnc*, encoding RNase III; and EM1332 [*rne*-*3071^ts^ zce*-*726*::Tn*10 rnc*::cam (gift of E. Masse and S. Gottesman)]. The *E. coli* strains were grown in LB containing 100 μg/mL ampicillin at 30° C until the OD_600_ reached ∼0.5; cultures were then switched to 42.5° C to inactivate RNase E as described (38).

### DNA and Proteins

The ampicillin resistant vector pBR-plac has been previously described (64). To generate pS17L, pS17S, and pGFK3038, S17 sequences from +1 to +188, from +124 to +188, and from +1 to +190, respectively, were cloned downstream of P*_lac_* between the AatII and EcoRI sites of pBR-plac (64) (Table S3; pS17L and pS17S were constructed by GenScript Biotech, Piscataway, NJ). pJN1 (65), which was used for *in vitro* transcription, contains the *B. pertussis* Tohama I sequence from 3363309 to 3363049 cloned between the EcoRI and HindIII sites in pTE103 (66). This cloning site places the +1 TSS of P_S17_ ∼365 nt upstream of the factor-independent, early transcription terminator of phage T7. The *in vitro* transcription plasmid p*fha*, which has been described (67), generates an RNA of ∼400 nt from P*_fhaB_*. Sequences of oligonucleotides used as probes and primers are available upon request. Sequences within plasmids were sequenced by Psomagen, Rockville, MD.

*B. pertussis* RNAP core, His_6_-tagged *B. pertussis* primary σ^A^, and BvgA were purified as described (19).

### RNA isolation, RNA-seq analyses and Northern blot analyses

*B. pertussis* Tohama I BP536 and its derivatives were grown on BG agar containing streptomycin (50 μg /ml) at 37° C in the absence [Bvg+, non-modulating mode] or presence [Bvg-, modulating mode] of 50 mM MgSO_4_ as described (25). After 3 days, cells were scraped from plates and resuspended in PLB medium (59) with or without MgSO_4_ to obtain an initial OD_600_ between 0.1 and 0.2. Cells were incubated at 37° C with shaking at 250 rpm until mid-log phase (OD_600_ ∼ 0.5). Culture aliquots of 7 ml were collected by centrifugation, total RNA was isolated by the hot-phenol method, and Northern analyses were performed as described (34). For RNA-seq analyses, total RNA was treated with DNase (Turbo DNA-free kit; Life Technologies, Inc.) for 1 h at 37° C, followed by phenol extraction and ethanol precipitation. After rRNA removal using the RiboZero kit (Illumina), strand-specific DNA libraries were prepared for Illumina sequencing using the ScriptSeq 2.0 kit (Illumina). Libraries were sequenced using a HiSeq 2500 sequencer (Illumina; NIDDK Genomic core Facility, single-end reads). Raw bulk RNA-seq data were processed using the following Snakemake (68) (v6.15.5) pipeline in the biowulf cluster: https://github.com/TriLab-bioinf/HINTON_LAB_TK_141/blob/main/Snakefile. Briefly, sequencing reads were quality trimmed and sequencing adapters were removed with fastp (v0.23.2). Trimmed reads were then mapped to the GenBank *B. pertussis* Tohama I assembly GCF_000195715.1 with STAR (v2.7.10a) and quantification of fragments per gene was carried out with the featureCounts tool from Subread software (v2.0.1) by counting reads falling within gene features. tRNAs and rRNAs were eliminated from our analysis. BigWig files were generated with bamCoverage tool from DeepTools (69) (v3.5.1). Processed bulk RNA-seq data were analyzed in R (v4.3.1) using the workflow in https://github.com/TriLab-bioinf/HINTON_LAB_TK_141/blob/main/TK_141_analysis.Rmd. Read counts were log-transformed with the DESeq2 function rlog. Differential gene expression analysis was performed separately for each dataset with the R package DESeq2 (70)(v1.42.0). Before the analysis, genes with less than 10 raw read counts in at least two samples were filtered out.

### RT-qPCR analyses

cDNA was generated using 2 μg of RNA, random hexamers (d(N)6; NEB; 1 nM) as primers and SuperScript IV Reverse Transcriptase (Thermo Fisher). RT-qPCR analyses were performed using a CFX96 Touch real-time detection system (NEB). Expression of the *B. pertussis* primary sigma factor gene (*rpoD*; *BP2184*) was used as an internal standard, and SsoAdvanced SYBR green Supermix (Bio-Rad) was used as the signal reporter. Reactions were performed in a 96-well microtiter PCR plate using 1 μl of cDNA at 2 ng/μl and sense and antisense primers (0.5 μM each) for amplifying each targeted gene in SsoAdvanced Universal SYBR green Supermix. Cycling conditions were as follows: denaturation (98° C for 30 s), amplification and quantification (95° C for 30 s, followed by 40 cycles of 95° C for 10 s, 55 to 60° C for 30 s, and 72° C for 30 s, with a single fluorescence measurement at 60° C for 30 s segments), a melting curve program (65 to 95° C with a heating rate of 0.5° C per second and continuous fluorescence measurement), and a cooling step to 65° C. Data were analyzed using the CFX manager software (Bio-Rad). The expression level of each sample was obtained by the standard curve for each gene and was normalized by the level of the internal control, primary sigma, *rpoD*. In each RT-qPCR analysis, the quantitation cycle (Cq_gene_) was observed for the gene of interest and for the reference gene (Cq*_rpoD_*); ΔCq = Cq_gene_- Cq*_rpoD_*; ΔΔCq = ΔCq for the ΔS17 strain - ΔCq for the WT strain; fold change (FC) = 2^-ΔΔCq^. For statistical analyses, the mean FC and standard error (SE) for a particular gene were determined using the website: https://ncalculators.com/statistics/standard-error-calculator.htm; the t statistic was calculated as the mean FC/SE, and the P value was determined using the two tailed hypothesis and a ‘degrees of freedom’ value of *N - 1* for the number of replicates (N) performed for a gene analyzed in a single RT-qPCR analysis and (*N_1_ - 1*) + (*N_2_ - 1*) + … for genes in which multiple analyses were performed using the website: https://www.socscistatistics.com/pvalues/tdistribution.aspx.

### 5’-RACE

5′-RACE was performed as described (71). Briefly, after DNase-treated total RNA was depleted of rRNA using the Ribo-Zero rRNA Depletion kit (Illumina), 1 μg (final volume 100 μL) of the resulting mRNA was incubated with and without RNA 5’-pyrophosphohydrolase (RppH; NEB, 50 U) at 37° C for 1 hour in the supplied 1X buffer (72), which generates 5’-monophosphorylated RNA from 5’-triphosphorylated mRNA. After extraction with 25:24:1 phenol/chloroform/isoamyl alcohol and precipitation with ethanol, the RNA was ligated to the 5′-universal RNA adapter (5′GAUAUGCGCGAAUUCCUGUAGAACGAACACUAGAAGAAA3′) using T4 RNA ligase (NEB). After extraction with 25:24:1 phenol/chloroform/isoamyl alcohol and ethanol precipitation, RNA (0.5 μg) was incubated with SuperScript IV reverse transcriptase (Thermo Fisher, Invitrogen), dNTPs, and Random Hexamers (Thermo Fisher) according to the manufacturer’s instructions. The resulting cDNA was then amplified by PCR using the P_BAD_ 5’- RACE universal primer (5’ACCTGACGCTTTTTATCGCAACTCTCTACTGTTTCTCCAT3’) and the S17-specific primer Ig*BP3151*-171R (5’CTTGCGATCTCGGGCTAAATCCACCA3’) in the presence of Q5 DNA polymerase (NEB) and dNTPs. PCR products were cloned using the Zero Blunt^TM^ TOPO^TM^ PCR cloning kit (Invitrogen), and the resulting plasmids were transformed into *E. coli* Top10 cells. Plasmids containing the amplified cDNA from the RNA with or without the RppH-treatment were sequenced using the M13For (−20) primer supplied with the TOPO cloning kit (Invitrogen).

### *In vitro* transcription

Transcription reactions were assembled as described (25) using 0.05 pmol of supercoiled DNA and as indicated, 5 pmol BvgA or BvgA∼P and 0.75 pmol *B. pertussis* RNAP (σ^A^:core at 2.5:1) in a total final volume of 10 μL. Where indicated, BvgA was phosphorylated by incubation in the presence of 20 mM lithium potassium acetyl phosphate (Ac∼P; Sigma-Aldrich) in 20 mM Tris-Cl, pH 8 for 30 min at room temperature immediately before use; nonphosphorylated BvgA was treated similarly without the addition of Ac∼P. Components were incubated at 37° C for 15 min before the initiation of a single round of transcription by the addition of heparin (final concentration of 200 μg/ml) and rNTPs (final concentration of 250 μM each ATP, GTP, and CTP and 25 μM [α-^32^P] UTP at 4.4 × 10^3^ dpm/pmol). Reaction products were separated on 4% (wt/vol) polyacrylamide, 7 M urea denaturing gels. After autoradiography, films were scanned using a Powerlook 2100XL densitometer.

### TEX (<u>T</u>erminator™ 5′-Phosphate Dependent <u>Ex</u>onuclease) digestion assay

Total RNA (7 ug), isolated with hot acid phenol extraction (34) from *B. pertussis* BP536 grown without or with MgSO_4_, was incubated with <u>T</u>erminator™ 5′-Phosphate Dependent <u>Ex</u>onuclease (TEX, Epicentre) or left untreated at 37° C for one hour as described (61). After electrophoresis on 10% TBE-UREA gels (Bio-Rad), RNA was transferred onto Zero probe GT blotting membranes (Bio-Rad) for Northern blot analysis. S17 and 5S RNA (as the control) were visualized with biotinylated probes.

### RNase assays

The *E. coli* strains EM1055, EM1277, EM1321, and EM1332, each harboring the S17 overexpressing plasmid pGFK3038, were grown in LB containing carbenicillin (100 μg/ml) at 30° C. At mid-log phase (OD_600_ of ∼0.5), cells were switched to 42.5°C and incubated for 15 min, which would inactivate RNase E in EM1277 and EM1332 (38). Aliquots (1 ml) of cell culture before and 15 min after the temperature switch were used to extract RNA with hot acid phenol as described (34). S17L and S17S transcripts were visualized by Northern blot. Biological triplicates of each sample were analyzed for statistical t-test.

### Rifampicin Chase

Three independent cultures of *B. pertussis* BP536 were grown in the absence of MgSO_4_ to an OD_600_ of ∼0.6. Rifampicin (Rif) was then added to a final concentration of 300 µg/ml, and 2 ml aliquots were taken at 0, 2, 4, 6, 8, 10, 20, and 60 min after Rif addition. Total RNA was extracted, and an aliquot (7 µg) was used for Northern analyses (as described above) to visualize the S17 RNA. The stable RNA SsrA was used as a control.

### β-Galactosidase Assays

*E. coli* strains, in which the *E. coli hfq* gene had been replaced with FLAG-tagged *B. pertussis hfq*, the indicated *B. pertussis* target-fused *lacZ* gene was present on the chromosome, and the vector (pBR-plac), pS17S, or pS17L was present, were streaked on LB agar plates containing 100 µg/ml of carbenicillin and incubated at 37° C overnight. The next day, 3-4 colonies were selected in combination and used to inoculate 3 mL of LB media containing 100 µg/ml of carbenicillin. After incubation overnight at 37° C with shaking, 10 mL LB containing 100 µg/ml carbenicillin, 0.02% arabinose, and 0.1 mM IPTG was inoculated to give an OD_600_ of ∼0.1, and cells were grown at 37 °C with shaking to an OD_600_ of 0.6-0.7. Miller assays were then performed essentially as described (73). In brief, cells were chilled on ice for 20 min to stop cell growth and a final OD_600_ was determined. A 500 µL aliquot was mixed with 500 µL Z buffer (60 mM Na_2_HPO_4_, 40 mM NaH_2_PO_4_, 10 mM KCl, 1 mM MgSO_4_, 50 mM Δ-mercaptoethanol, pH adjusted to 7 using 8 M NaOH), 20 µL 0.1% SDS, and 40 µL chloroform. After incubation at room temperature for 15 min, 500 µL of the aqueous phase was transferred to a new tube and 100 µL 4 mg/ml *ortho*- Nitrophenyl-β-galactoside (ONPG) was added to initiate the β-galactosidase reaction. After the development of a yellow color, reactions were neutralized by the addition of 250 µL 1 M Na_2_CO_3_. The OD_550_ and OD_420_ were measured using a blank (250 µL LB media, 250 µL Z buffer, 100 µL ONPG, and 250 µL Na_2_CO_3_). β-galactosidase units (Miller units) were determined using the equation:

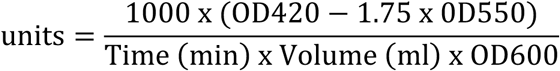

Experiments were conducted in biological duplicates and technical triplicates, and relative means and one-way ANOVA tests to determine statistical significance were performed in Prism 10 for Mac.

### Growth of *E. coli* containing an overexpression S17 plasmid

BL21 containing pBR-plac was grown in LB broth plus 100 μg/ml carbenicillin at 37° C with shaking to an OD_600_ of ∼0.3. Cultures were split, water (noninduced) or IPTG (final concentration of 1 mM) was added, and continued growth was monitored by following OD_600_.

## ACKNOWLEDGEMENT

We thank Susan Gottesman and Eric Masse (National Cancer Institute) for providing the *rne* and *rnc* mutant strains and Brooke Weingard (NIDDK, National Institutes of Health) for technical help with the tables. This work utilized the computational resources of the NIH HPC Biowulf cluster.

