## Supplementary Figures for "A highly conserved sRNA downregulates multiple genes, including a σ^54^ transcriptional activator, in the virulence mode of *Bordetella pertussis*"

### Supplemental Figures

Fig. S1

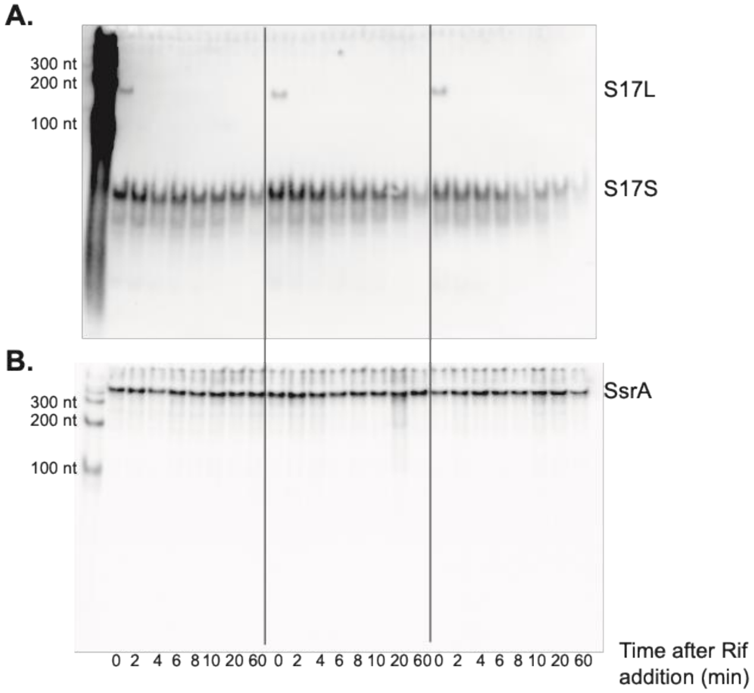


FIG S1 S17S is more stable than S17L *in vivo*. Northern blots showing the levels of S17 RNA (top) or SsrA (bottom, same blot after rehybridization) in 3 biological replicates of WT samples, grown without $MgSO_4$, before (0 time) and after the addition of rifampicin (Rif) for the indicated times. Lane 1 shows a size marker ladder with the positions of 100, 200, and 300 nucleotides (nt) indicated. The positions of S17L, S17S, and SsrA are indicated.

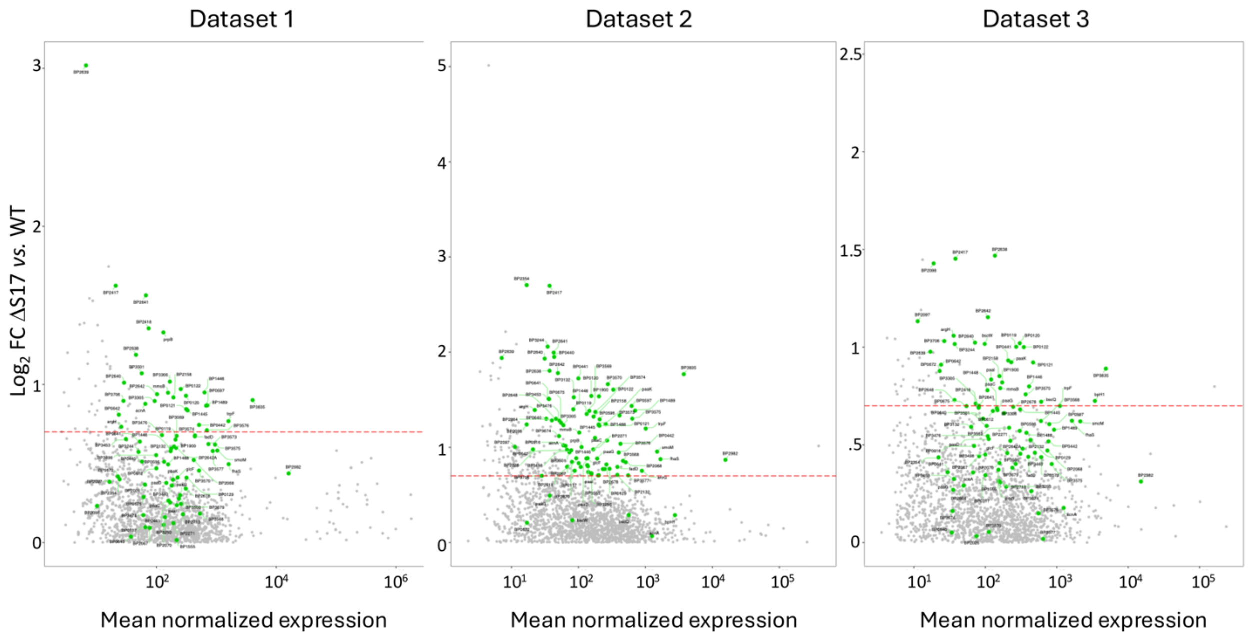


FIG S2  Plots showing the log2FC for the ΔS17 vs. WT strains from RNA-seq datasets 1, 2, and 3 *vs.* the mean normalized expression.  Green dots indicate any gene that had a log2FC > 0.7 with an adjusted p-value of < 0.05 in any one of the datasets.  Grey dots show all the other genes with log2FC > 0.

FIG. S3

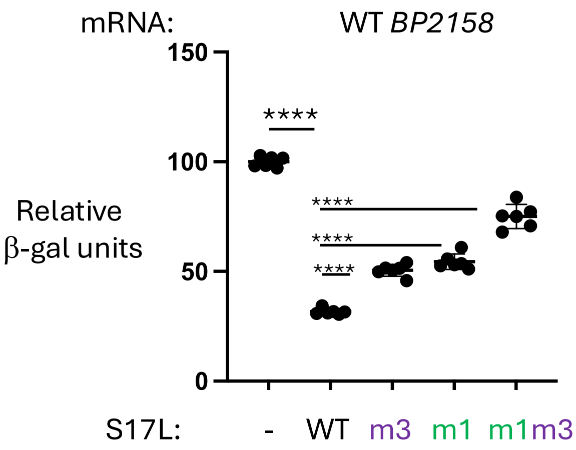


FIG S3  S17L post-transcriptionally represses *BP2158*.  Results of β-galactosidase (β-gal) assays showing Miller units, using the strain with WT *BP2158* and the indicated S17L plasmids, relative to a plasmid without an S17L insert (-).  Means and standard deviations are indicated by the horizontal lines among the data points.  In some cases, the points are too close together to be individually distinguishable.  Results of one-way ANOVA comparison tests for various datasets are indicated: ****, p-value < .0001.

FIG. S4

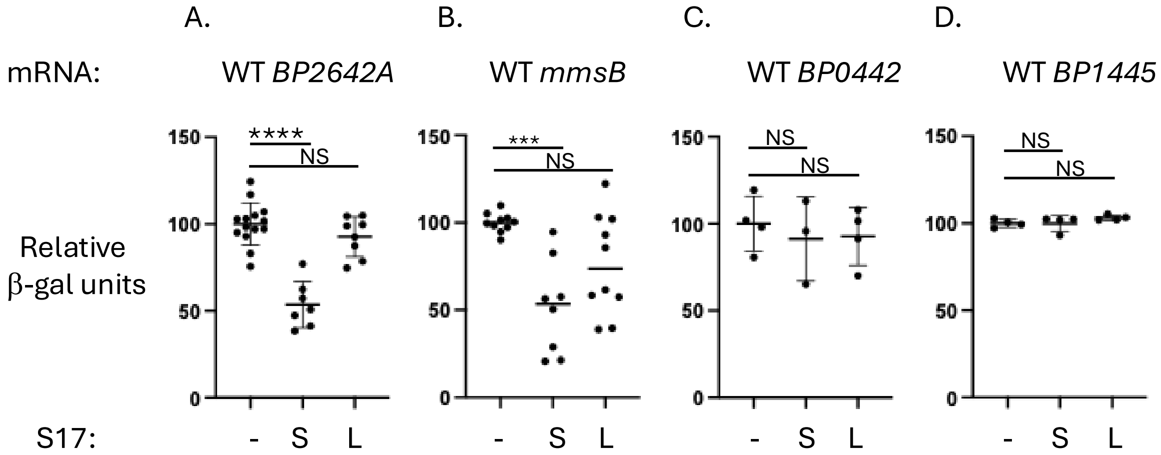


FIG S4 S17S post-transcriptionally represses *BP2642A* and *mmsB*. Results of β-galactosidase (β-gal) assays showing the Miller units obtained with the following strains containing the indicated S17S or S17L plasmids, relative to one containing the plasmid without a S17 insert (-): (A) WT *BP2642A*, (B) WT *mmsB*, (C) WT *BP0442*, and (D) *BP1445*. Means and standard deviations are indicated by the horizontal lines among the data points. In some cases, the points are too close together to be individually distinguishable. Results of one-way ANOVA comparison tests for various datasets are indicated: NS, not significant; ***, p-value < .001; ****, p-value < .0001.

FIG. S5

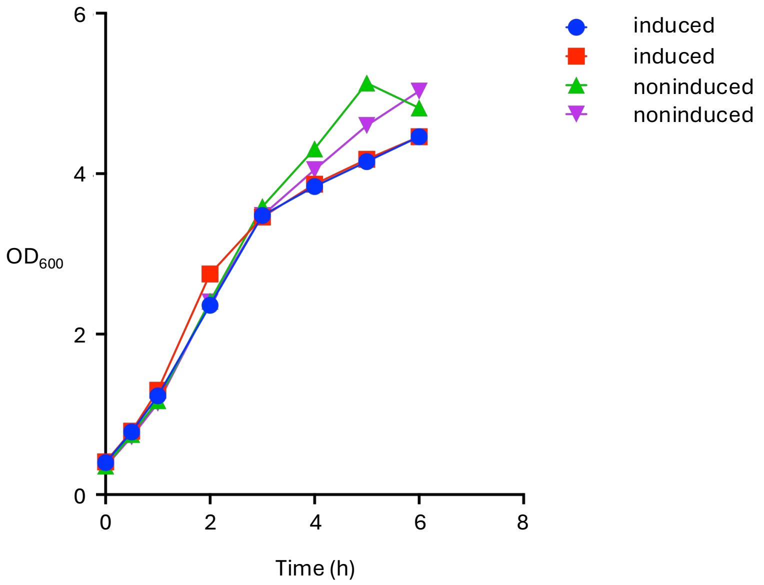


FIG S5 Overexpression of S17 does not inhibit *E. coli* growth at 37$^{o}$ C. Growth (measured by $OD_{600}$) of 2 independent cultures of BL21 containing pS17S after the addition of 1 mM IPTG (blue circles and orange squares) or an equivalent volume of water (green and purple triangles). Plot is representative of 2 experiments.

**Supplemental Tables**

Table S1. RNA-seq differential analyses of ΔS17 *vs.* WT for dataset 1 (sheet 1), 2 (sheet 2), and 3 (sheet 3).

Table S2. Analyses of significantly up-regulated genes in the absence of S17. Sheet 1, genes identified as significantly up-regulated; sheet 2, correlation between genes up-regulated in the absence of S17 and genes found in the Δ*hfq vs.* WT dataset (12); sheet 3, homologs of various significantly up-regulated genes found in representative Betaproteobacteria and their possible S17 binding sites; sheet 4, genes significantly down-regulated by the *B. cenocepacia* sRNA *bdhR1* (42).

Table S3. Sequences of DNA inserts
